# Microbial abundance and diversity in 64-74 Ma subseafloor igneous basement from the Louisville Seamount Chain

**DOI:** 10.1101/2023.11.02.565174

**Authors:** Jason B. Sylvan, Benjamin J. Tully, Yuki Morono, Jeffrey C Alt, Sharon L. Grim, Fumio Inagaki, Anthony A.P. Koppers, Katrina J. Edwards

**Affiliations:** Department of Biological Sciences, University of Southern California, Los Angeles, CA 90089, USA; Center for Dark Energy Biosphere Investigations, University of Southern California, Los Angeles, CA 90089, USA; Kochi Institute for Core Sample Research, Japan Agency for Earth-Marine Science and Technology (JAMSTEC), Nankoku, Kochi 783-8502, Japan; Department of Earth and Environmental Sciences, University of Michigan, Ann Arbor, MI 48109, USA; Josephine Bay Paul Center for Comparative Molecular Biology and Evolution, Marine Biological Laboratory, Woods Hole, MA 02543, USA; Advanced Institute for Marine Ecosystem Change (WPI-AIMEC), JAMSTEC, Yokohama 236-00001, Japan; Department of Earth Sciences, Graduate School of Science, Tohoku University, Sendai 980-8574, Japan; College of Earth, Ocean and Atmospheric Sciences, Oregon State University, Corvallis, Oregon 97331, USA; Department of Oceanography, Texas A&M University, College Station, TX 77845, USA

## Abstract

The aquifer in subseafloor igneous basement is a massive, continuous microbial substrate, yet sparingly little is known about life in this habitat. The work to date has focused largely on describing microbial diversity in young basement (<10 Ma) at oceanic spreading regions and ridge flanks, where the basaltic crust is still porous and fluid flow through it is active. While the prevailing belief used to be that fluid flow through older parts of the seafloor was non-existent, recent heat flow models predict that fluid moves through subseafloor basement >65 Ma, and that seamounts can act as mid-plate conduits for fluids into and out of the subsurface aquifer in older crustal settings. Here we test the hypothesis that microbial life exists in subseafloor basement >65Ma using samples collected from the Louisville Seamount Chain via seafloor drilling. Cell biomass was heterogeneous in nature and ranged from below detection to ∼10^4^ cells cm^-3^. Bacterial 16S rRNA genes from core samples and enrichment incubations are dominated by lineages putatively carrying out hydrocarbon oxidation and nitrogen, sulfur and metal redox processes. Samples from two different seamounts were statistically different, indicating some degree of biogeography. Archaea were not detected via quantitative polymerase chain reaction, indicating they are rare in the Louisville subsurface. Taken together, the data indicate that microbial life is indeed present in subseafloor igneous basement >65 Ma, which significantly expands the range of the subseafloor biosphere where microbial life is known to exist.

**Impact Statement:** The aquifer in subseafloor igneous basement is the largest continuous microbial substrate on Earth, but it is difficult to access and therefore understudied. We here collected samples from the Louisville Seamount Chain using seafloor drilling to determine if microbial life exists in the >65 Ma subseafloor basement made at these seamounts. A low biomass environment dominated by Bacteria potentially capable of using the Fe and S inherent in subseafloor basalt was detected, including Bacteria that were revived in enrichment experiments. This discovery expands the range of seafloor where confirmed microbial life exists and indicates the interior of seamounts is habitable.

## Introduction

The entire volume of the ocean cycles through subseafloor basaltic crust every 10^5^-10^6^ years^1^, and the volume of the habitable zone in marine basement is equivalent to the entire volume of the marine water column^2,3^. The upper oceanic crust, represented by basaltic rock, is therefore an important microbial substrate. Recent work investigating microbial communities on basalts exposed at the seafloor revealed that resident microbial communities are abundant, diverse and highly active^4–7^. Despite the potential importance and magnitude of the microbial biosphere in subseafloor basalts, knowledge about this biosphere remains extremely limited, partially due to the difficulty of sampling this environment^8^. To date, only a few studies have focused on microbial processes in subseafloor basalts, and primarily targeting young basalts <10 Ma. For example, it was recently shown that microbial communities in deeply buried fluids moving through 3.5 Ma basalts on the Juan de Fuca Ridge (JdFR) are annually variable and taxonomically and functionally consistent with the warm, anoxic fluids there^9–12^. Cell abundances from these boreholes are ∼10^4^ cells ml^-1^, indicating low but measurable biomass in this young, hydrologically active environment^12^. Analysis of biofilms growing on rocks in incubations in the JdFR boreholes and *in situ* core samples revealed the presence of both sulfate reducing and methanogenic microorganisms^13–16^.

Most recently, a series of experiments resulting from drilling and installing multiple borehole observatories in a cool, ridge-flank environment adjacent to the Mid-Atlantic Ridge, called North Pond^17^, has resulted in a series of breakthroughs yielding insight into microbial life in subseafloor basaltic basement. Fluids circulating through basaltic basement at North Pond are cool (∼4°C) and oxic, in stark contrast to JdFR^18^. Subsurface fluids in North Pond are oxygenated and provide a source of re-oxygenation of deep sediments in the sediment pond^19^ - modeling of oxygen flux indicated that microbes in the basaltic crust consume oxygen at a rate of ∼1 nmol cm^-3^ day^-1 20^. Microbial respiration rates for both heterotrophy and autotrophy in basement were measured to be higher than deep seawater just above the seafloor at the site^21^, and multi-year sampling revealed the capacity for hydrogen, sulfur and iron oxidation amongst a microbial community that shifted over time while retaining similar functional capabilities^22,23^. Analysis of cored rocks from North Pond revealed cell densities from below detection to ∼6x10^4^ cells cm^-3 24^, compared to cell densities in the fluids of 5.1x10^3^ - 2.8x10^4^ cells ml^-1 21,23^. The cored rocks also revealed the presence of a microbial community dominated by Proteobacteria, while Archaea are present at relative abundances of <1%^25^.

Analysis of deep subseafloor microbial communities in lower oceanic crust, comprised largely of gabbro, accessed at sites where tectonic windows bring it close to the seafloor, reveal further insight into subseafloor basement microbial community diversity and function. Microbial communities detected up to 1391 m below seafloor (mbsf) in Hole U1309D, comprised of lower oceanic crust exposed at the 1.5 Ma Atlantis Massif, suggest the presence of an ecosystem where Archaea are rare and hydrocarbon degradation appears to be a common metabolism ^26^. Newly developed cell enumeration protocols revealed cell abundance from below detection up to 1.6x10^4^ cells cm^-3^ in samples from seven boreholes drilled in Altantis Massif that cored up to 15 mbsf ^27^. Cell densities for the majority of samples, however, was <10^3^ cells cm^-3^. At Atlantis Bank, an oceanic core complex on the Southwest Indian Ridge, cell counts in Hole U1473A from 10-781 mbsf were also ∼10^4^ cells cm^-3^ or less, and microbial diversity and abundance was variable downcore ^28^.

The pioneering work done on the JdFR, North Pond, Atlantis Massif and Atlantis Bank all indicate heterogeneous microbial communities in young subseafloor basement. Efforts to understand microbial life in older settings are even more scarce and have been largely hampered by low biomass recovery and drilling contamination^8,29^. As a global average, oceanic hydrothermal circulation is predicted to cease at 65 Ma due to crustal sealing^30–32^, so interest has been higher in younger crust with potentially high rates of fluid flow. However, there is indeed evidence for fluid flow in older crust^33–35^, suggesting that a subseafloor biosphere in igneous basement may also exist there. In fact, recent evidence from basement underlying the South Pacific Gyre revealed locally high cell densities at grain boundaries, potentially up to 10^10^ cells cm^-3^ in samples from 33.5 and 104 Ma basalt^36^, but this tantalizing evidence was restricted to two samples.

It is known that seamounts provide conduits of fluid flow into and out of basement^37,38^, and that this is true even in basement >65 Ma^39^, indicating that seamounts located on older parts of the ocean crust may be home to subsurface microbial populations carried there by entrained seawater or resident since sometime after eruptive activity created the seamount. To test the hypothesis that subseafloor igneous rocks in seamounts >65 Ma host microbial populations, and to expand our understanding about life in basement in the understudied but most expansive biome on earth, we collected samples cored during seafloor drilling into extinct seamounts along the Louisville Seamount Chain during Integrated Ocean Drilling Program (IODP) Expedition 330^40^. Drilled rock cores were collected from 29-491 mbsf from Holes U1372A (74.2 Ma), U1373A (69.5 Ma), U1374A (67.4-71.1 Ma) and U1376A (64.1 Ma (Figure 1), all consisting of basement created at the Louisville hotspot in the south Pacific Ocean and described as alternating units of sediment or volcaniclastic breccia and erupted basalt^40^. Here we report results from those samples for cell enumeration, measurement of organic carbon content, δ^13^C stable isotope analysis, and 16S rRNA amplicon analysis of rock core samples and enrichment cultures.

**Figure 1.**
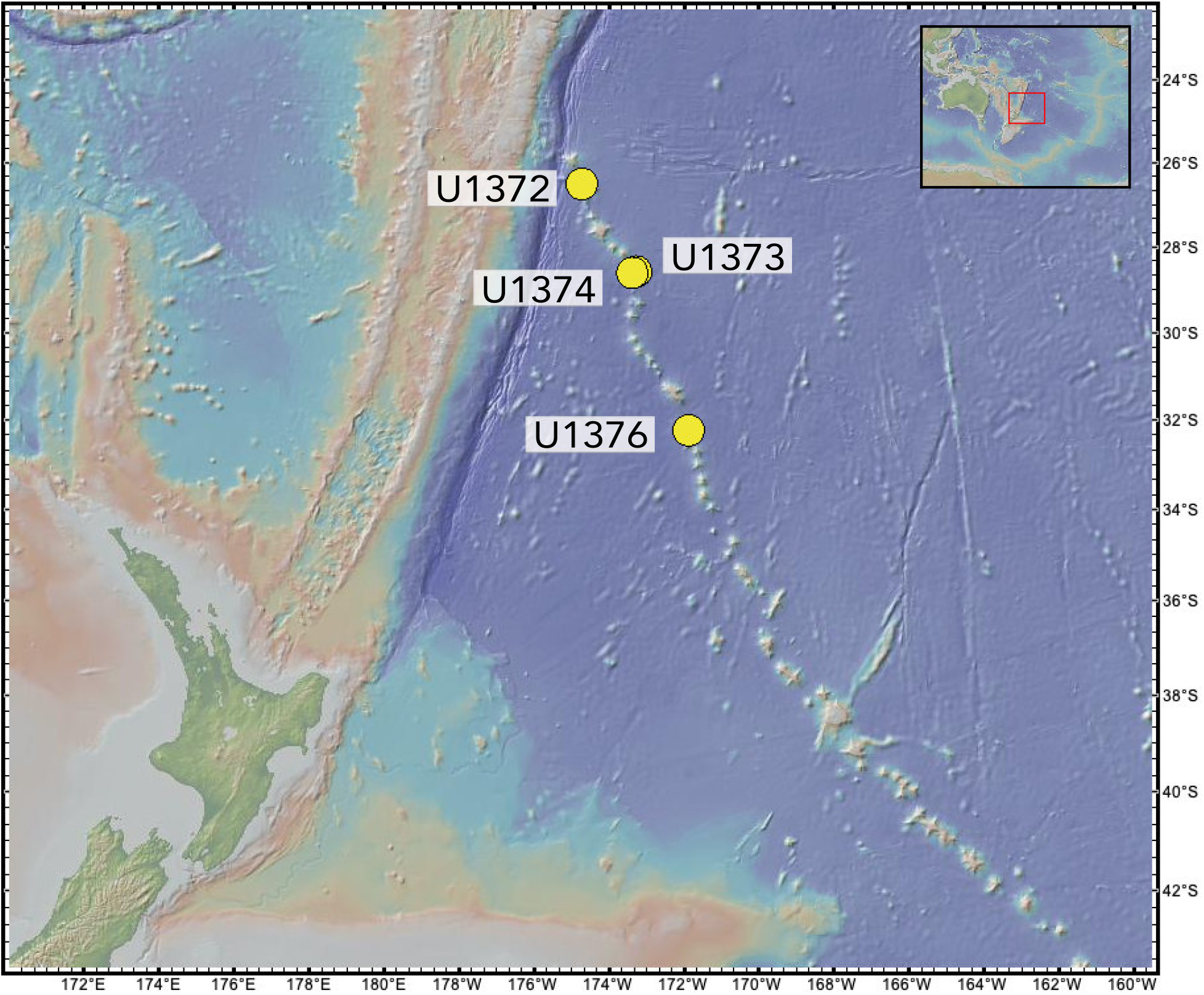
Map of Sites drilled and cored during IODP Expedition 330 from which samples were analyzed for this project. Figure made with GeoMapApp (www.geomapapp.org) / <u>CC BY</u> / CC BY (Ryan et al., 2009)^94^.

## Results/Discussion

We quantified cell density in 53 samples from the Louisville Seamounts, including 13 from Canopus Guyot (Site U1372), 29 from Rigil Guyot (Site U1374, including one sample from sediment overlying igneous basement), and 11 from Burton Guyot (Site U1376). We also analyzed 6 samples from Site U1301B on the JdFR flank for context^13^. Prokaryotic cell abundance in our samples ranges from below detection (no cells detected on the filter) to 8.3x10^4^ cells cm^-3^ with a minimum quantifiable limit of 1.20x10^2^ cells cm^-3^ (Figure 2). Cell concentrations from the young JdFR flank are similar to those from the Louisville Seamounts, and the single sediment sample falls within the expected range for samples from the South Pacific Gyre ^41^, in which Site U1374 is located. There is no apparent relationship between lithology or depth and biomass, unlike subsurface biomass in sedimentary environments, where biomass decreases from the seafloor in a linear fashion on a log scale ^42^.

**Figure 2.**
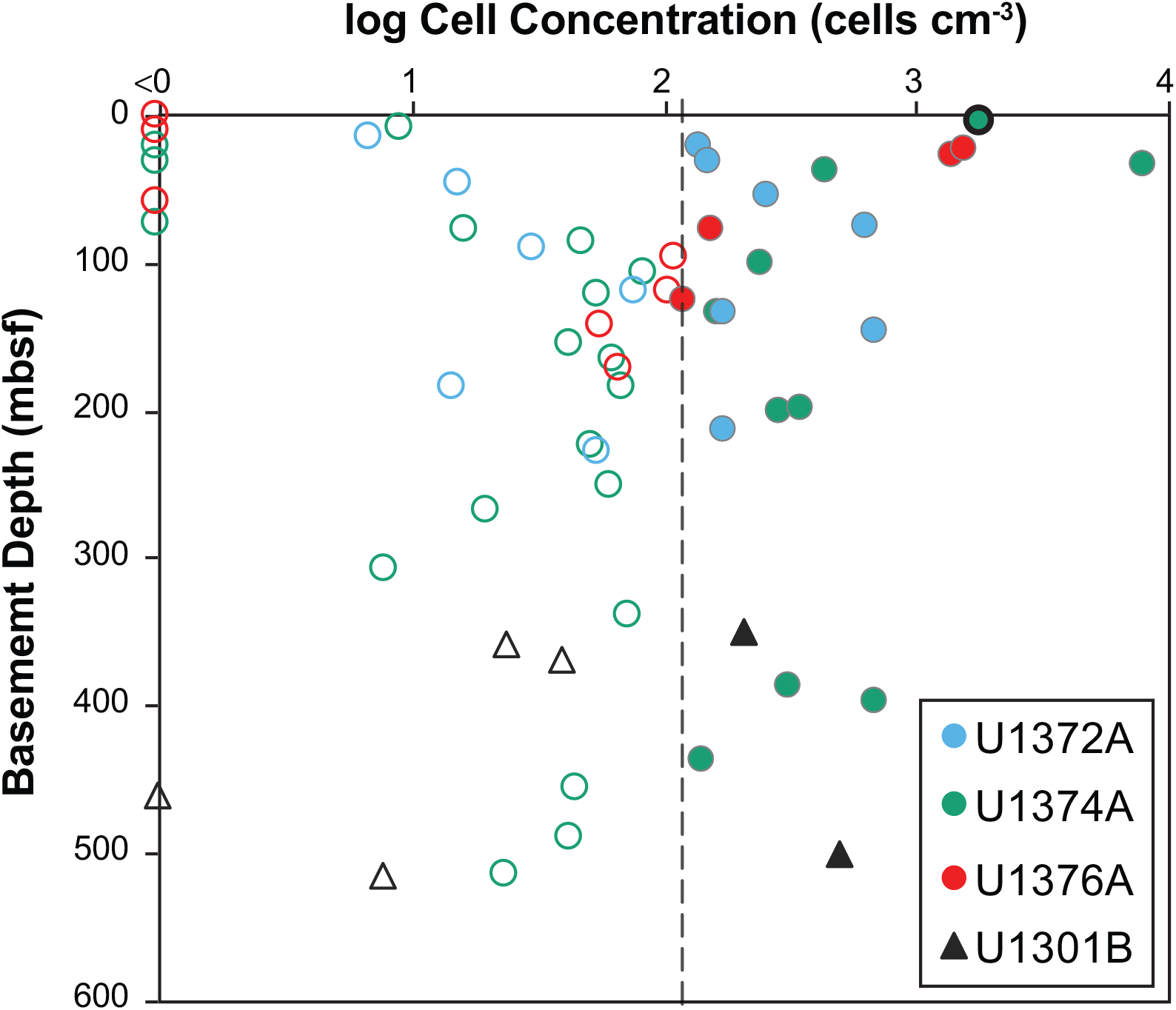
Subseafloor cell counts from IODP Expeditions to the Louisville Seamounts (Expedition 330, U1372A, U1374A and U1376A) and the Juan de Fuca Ridge flank (Expedition 301, U1301B). The dotted line indicates the limit of quantification, values to the left are considered below quantification. The sample from Site U1374A circled in black represents a sediment sample overlying basement at that site.

The cell concentrations detected here are similar in magnitude to deep subseafloor basement in the JdFR, evidenced by whole-round core samples analyzed here as well as observatory fluid samples from Site U1362, which range 2.6x10^3^ - 2.6x10^4^ cells cm^-3 11^. Similar concentration ranges were also detected in the cool, oxic ridge flank environment near the Mid-Atlantic Ridge at the North Pond study site, where samples from whole-round cores had cell concentrations ranging below detection to 6x10^4^ cells cm^-3 24^ and fluid samples from borehole observatories ranged 1-2x10^4^ cells cm^-3 21^, as well as in upper oceanic crust from 10-781 mbsf at the 12 Ma Atlantis Bank^28^. Cell densities in shallow Atlantis Massif, ∼1.5 Ma, are the lowest of the basement sites studied to date^27^. Of particular interest, cell concentrations in the whole-rock samples from North Pond, Atlantis Bank and Atlantis Massif, are also very heterogeneous. Data from the Louisville Seamount Chain and JdFR presented here, coupled with the data from these other basement sites, add the growing consensus that heterogeneous abundance with depth is likely a defining characteristic of subseafloor basement microbial communities. This indicates that there are fundamental differences between controls on subsurface life in marine basement versus sediments. We hypothesize that this is driven by fluid flow pathways in basement, which are heterogeneous, whereas biomass distribution in sediments is largely constrained by biomass in overlying waters ^42^. Therefore, predictions of global biomass in the subseafloor aquifer must account for fracture density and porosity and permeability structure in basement.

Weight % total C, total organic carbon (TOC), and 8^13^C for total C and TOC were measured from a subset of the samples analyzed for biomass that had enough material (n=8). Weight % TOC ranges from 0.006 to 0.154 (Table S1), which is generally lower than subseafloor basalts collected from Hole U1301B^13^. No relationship is evident between microbial cell counts and weight % TOC, suggesting that microbial biomass makes up a small percentage of TOC in the Louisville Seamounts basement. 8^13^C-TOC values are -17.8 to -23.7‰, which is slightly higher than for subseafloor basalts from JdFR, -21.6 to -34.6‰, which were interpreted to reflect the presence of microbial organic carbon in the rocks^13^. The 8^13^C-TOC values for the Louisville seamounts are generally similar to that for dissolved organic carbon in seawater (813C = -20 to -22‰; Druffel et al., 1992), and the simplest explanation is that seawater is the source, with no fractionation occurring in the subsurface. An alternative hypothesis is that the TOC is a product of microbial activity in the rocks with seawater carbonate or DOC as the source, which is supported by indications of recycling of organic matter at Site U1473 on Atlantis Bank ^43^. The slightly more negative 8^13^C-TOC values (to -23.7‰) could be the result of microbial activity, similar to that at JdFR^13,44^. Some samples from borehole U1301B on the JdFR are heterogeneous, with some similar to seawater DOC, while others are in the -27 to -34‰ range^13^, indicating that microbial effects in basement rocks are variable, depending on fluid flow pathways in the basement, as well as heterogeneity of alteration effects.

We used amplicon sequencing of the V4V6 region of the bacterial 16S rRNA gene^45^ to examine bacterial diversity in 15 samples from Holes U1374A (9 samples, 40-491 mbsf) and U1376A (6 samples, 29-174 mbsf (Table 1)). Note that short sample names used in the text and figures are the last two numbers of the site followed by the core number (e.g. 7407 for sample U1374A-7R1). Almost no archaeal 16S rRNA genes were detected via qPCR for all of these samples except for 1376-23, indicating Archaea are either absent or present at very low abundance in this environment (see discussion in SOM for further details). Similar lack of archaeal biomass was detected in subsurface basalt samples from North Pond^25^ and gabbroic samples from Atlantis Bank^28,43^ and is common for basalts exposed at the seafloor^7,46,47^. This is not the case at the JdFR sites^11–13^, which are warm (∼65°C) and anoxic. Altogether, these findings support the trend that Archaea have low abundance on cold silicate rocks.

**Table 1.**
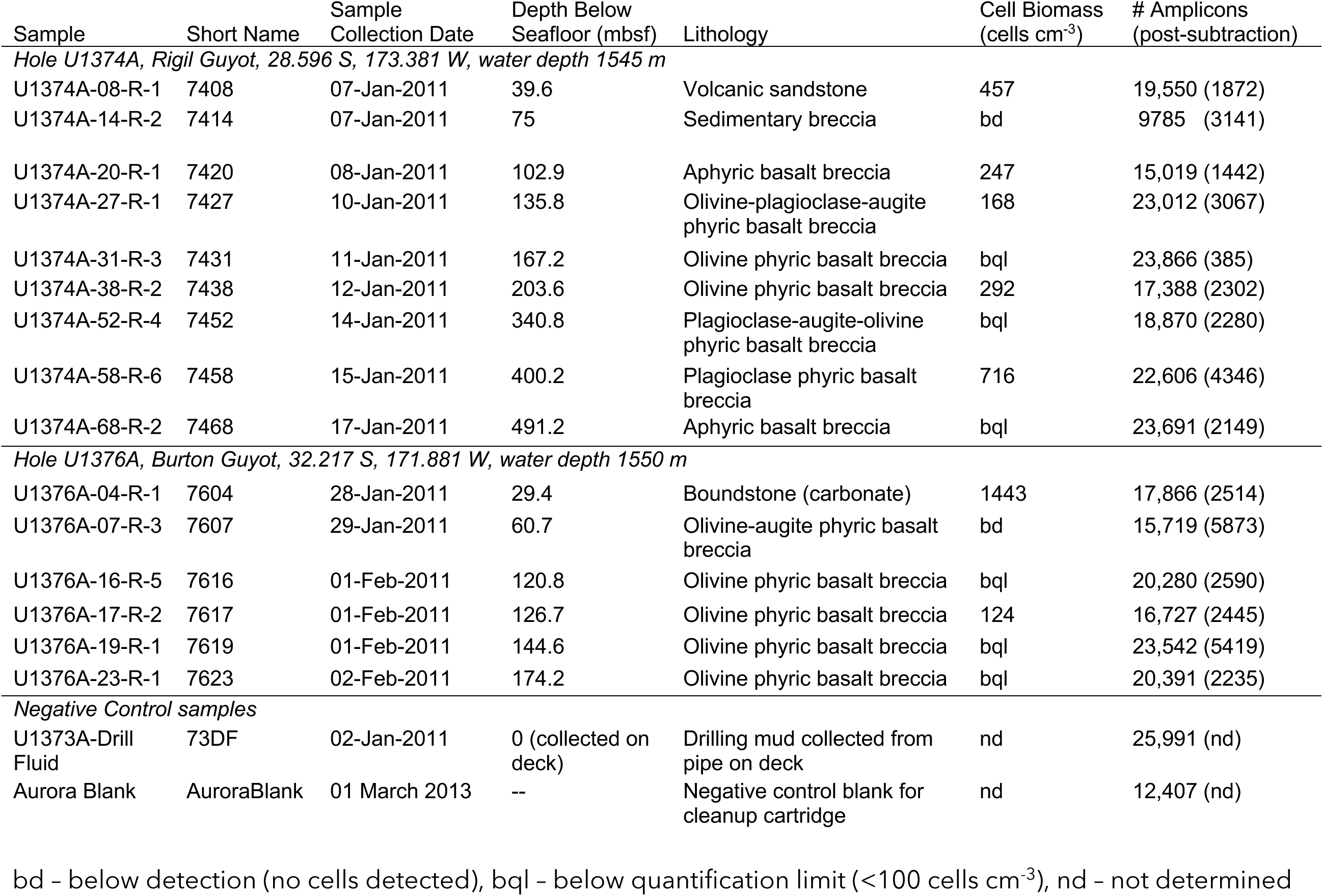
Sample descriptions and cell biomass. Short sample names are the last two numbers of the site followed by the core number (e.g. 7407 for sample U1374A-7R1).

| Sample | Short Name | Sample<br>Collection Date | Depth Below<br>Seafloor (mbsf) | Lithology | Cell Biomass<br>(cells cm<sup>-3</sup>) | # Amplicons<br>(post-subtraction) |
| --- | --- | --- | --- | --- | --- | --- |
| *Hole U1374A, Rigil Guyot, 28.596 S, 173.381 W, water depth 1545 m* |  |  |  |  |  |  |
| U1374A-08-R-1 | 7408 | 07-Jan-2011 | 39.6 | Volcanic sandstone | 457 | 19,550 (1872) |
| U1374A-14-R-2 | 7414 | 07-Jan-2011 | 75 | Sedimentary breccia | bd | 9785 (3141) |
| U1374A-20-R-1 | 7420 | 08-Jan-2011 | 102.9 | Aphyric basalt breccia | 247 | 15,019 (1442) |
| U1374A-27-R-1 | 7427 | 10-Jan-2011 | 135.8 | Olivine-plagioclase-augite<br>phyric basalt breccia | 168 | 23,012 (3067) |
| U1374A-31-R-3 | 7431 | 11-Jan-2011 | 167.2 | Olivine phyric basalt breccia | bql | 23,866 (385) |
| U1374A-38-R-2 | 7438 | 12-Jan-2011 | 203.6 | Olivine phyric basalt breccia | 292 | 17,388 (2302) |
| U1374A-52-R-4 | 7452 | 14-Jan-2011 | 340.8 | Plagioclase-augite-olivine<br>phyric basalt breccia | bql | 18,870 (2280) |
| U1374A-58-R-6 | 7458 | 15-Jan-2011 | 400.2 | Plagioclase phyric basalt<br>breccia | 716 | 22,606 (4346) |
| U1374A-68-R-2 | 7468 | 17-Jan-2011 | 491.2 | Aphyric basalt breccia | bql | 23,691 (2149) |
| *Hole U1376A, Burton Guyot, 32.217 S, 171.881 W, water depth 1550 m* |  |  |  |  |  |  |
| U1376A-04-R-1 | 7604 | 28-Jan-2011 | 29.4 | Boundstone (carbonate) | 1443 | 17,866 (2514) |
| U1376A-07-R-3 | 7607 | 29-Jan-2011 | 60.7 | Olivine-augite phyric basalt<br>breccia | bd | 15,719 (5873) |
| U1376A-16-R-5 | 7616 | 01-Feb-2011 | 120.8 | Olivine phyric basalt breccia | bql | 20,280 (2590) |
| U1376A-17-R-2 | 7617 | 01-Feb-2011 | 126.7 | Olivine phyric basalt breccia | 124 | 16,727 (2445) |
| U1376A-19-R-1 | 7619 | 01-Feb-2011 | 144.6 | Olivine phyric basalt breccia | bql | 23,542 (5419) |
| U1376A-23-R-1 | 7623 | 02-Feb-2011 | 174.2 | Olivine phyric basalt breccia | bql | 20,391 (2235) |
| *Negative Control samples* |  |  |  |  |  |  |
| U1373A-Drill<br>Fluid | 73DF | 02-Jan-2011 | 0 (collected on<br>deck) | Drilling mud collected from<br>pipe on deck | nd | 25,991 (nd) |
| Aurora Blank | AuroraBlank | 01 March 2013 | -- | Negative control blank for<br>cleanup cartridge | nd | 12,407 (nd) |

bd – below detection (no cells detected), bql – below quantification limit ( $<100 \text{ cells cm}^{-3}$ ), nd – not determined

To control for potential contamination of our samples by drilling fluid or during DNA extraction and cleanup, we first generated OTUs at the 95% similarity level and subtracted OTUs detected in negative controls at abundances 10X higher than experimental samples. The 10X threshold was used to allow for the possibility that some OTUs may show up in the controls but be much more abundant in the samples, indicating they are likely true members of the community in that sample. This approach avoids overly conservative discarding of any sequence detected in the controls but is still strict enough to minimize the likelihood of contaminant sequences remaining after QC. After removing contaminants detected at the 95% OTU level, we reanalyzed the sequences that passed QC by constructing OTUs at 97% similarity cutoff. Following this quality control step, there were 385-5873 bacterial V4V6 amplicons per sample (Table 1). The percent of amplicons remaining following blank subtraction ranged ∼2-37% with a mean value of 16%.

Mean bacterial communities sampled from Hole U1376A were more diverse than those from Hole U1374A (Table S2, Figure S1), but this was only statistically significant for richness (t-test, p<0.023). The most abundant phyla represented in the subseafloor Louisville Seamount environment are the Actinobacteriota, Bacteroidota, Firmicutes and Proteobacteria (Figure S2). Within those phyla, the classes Actinobacteria, Bacteroidia, Clostridia, Alpha-, and Gammaproteobacteria are the most abundant.

For those top five most abundant classes, one to six orders per lineage represented the majority of diversity present (Figure 3). Amongst these, Corynebacterales, Bacillales, Clostridiales, Rhizobiales, Burkholderiales and Xanthomonadales were the most abundant. Burkholderiales were also abundant community members in subseafloor basalts underlying North Pond^25^. Actinobacteria and Firmicutes were found to be more abundant at U1374A and U1376A than at the North Pond holes.

**Figure 3.**
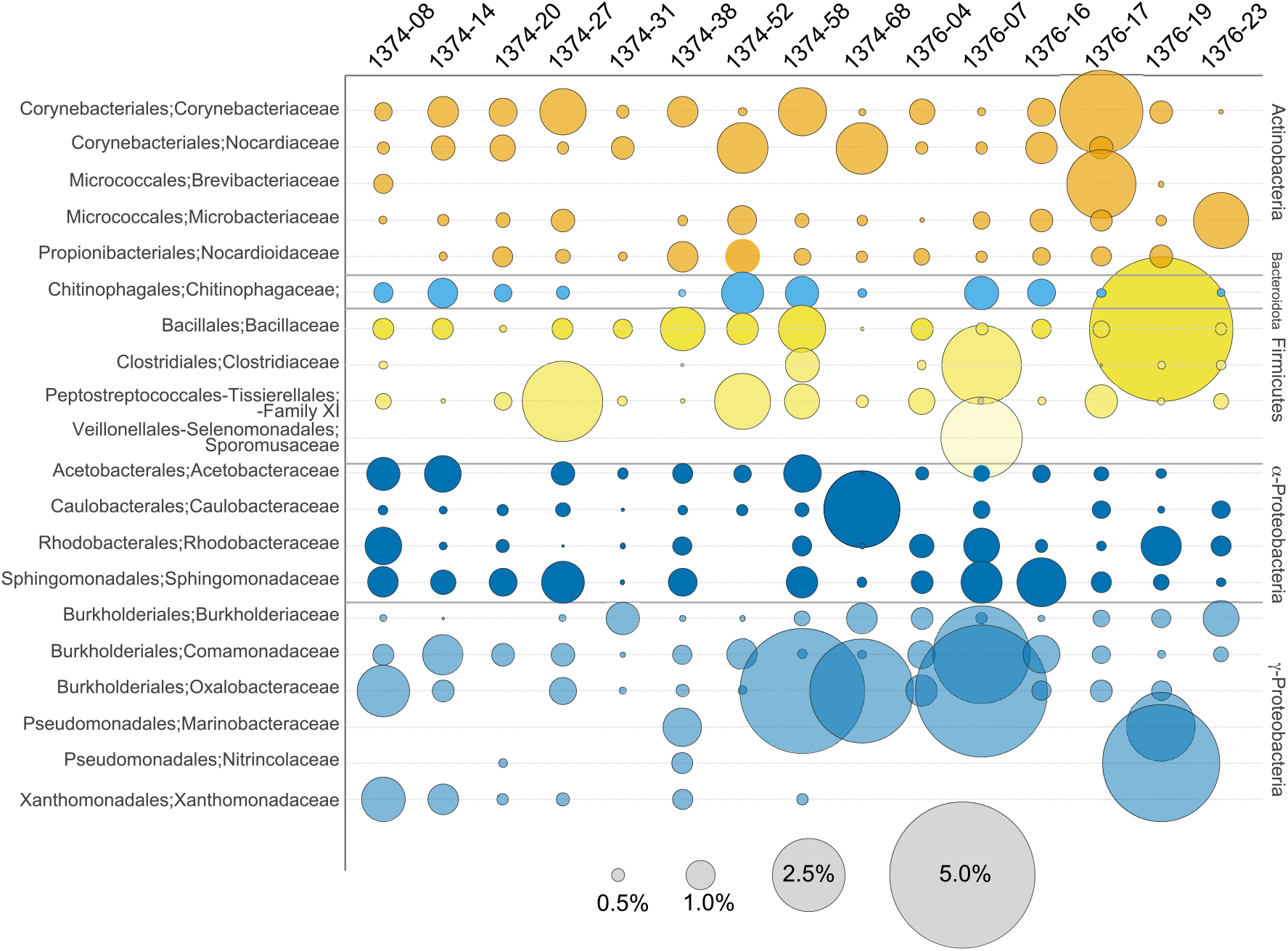
Relative abundance of bacterial families representing ≥1% of the total dataset. Order and family names are on the left, Phylum and class (for Proteobacteria) are on the right. Color code matches class in Figure S2.

Analysis of community richness reveals that microbial communities in samples from Holes U1374A and U1376A are most similar to other samples from the same Hole (Figure 4). More specifically, two clusters are apparent: microbial communities from samples 1374-08, 1374-14, 1374-20, 1374-27 and 1374-38, all from Hole U1374A, grouped together, while communities from samples 1376-04, 1376-16 and 1376-19, from Hole U1376A, formed a separate group. This indicates that location and/or seamount has some impact on community structure in this environment. Importantly, there were no apparent patterns between microbial communities and lithology, indicating that the individual seamount, and therefore Site location, or perhaps age of substrate (since age varies by seamount), may have a more important impact in this case than lithology. Prior work with basalts at the seafloor revealed that community similarity is strongly influenced by age of the rock sampled^5^.

**Figure 4.**
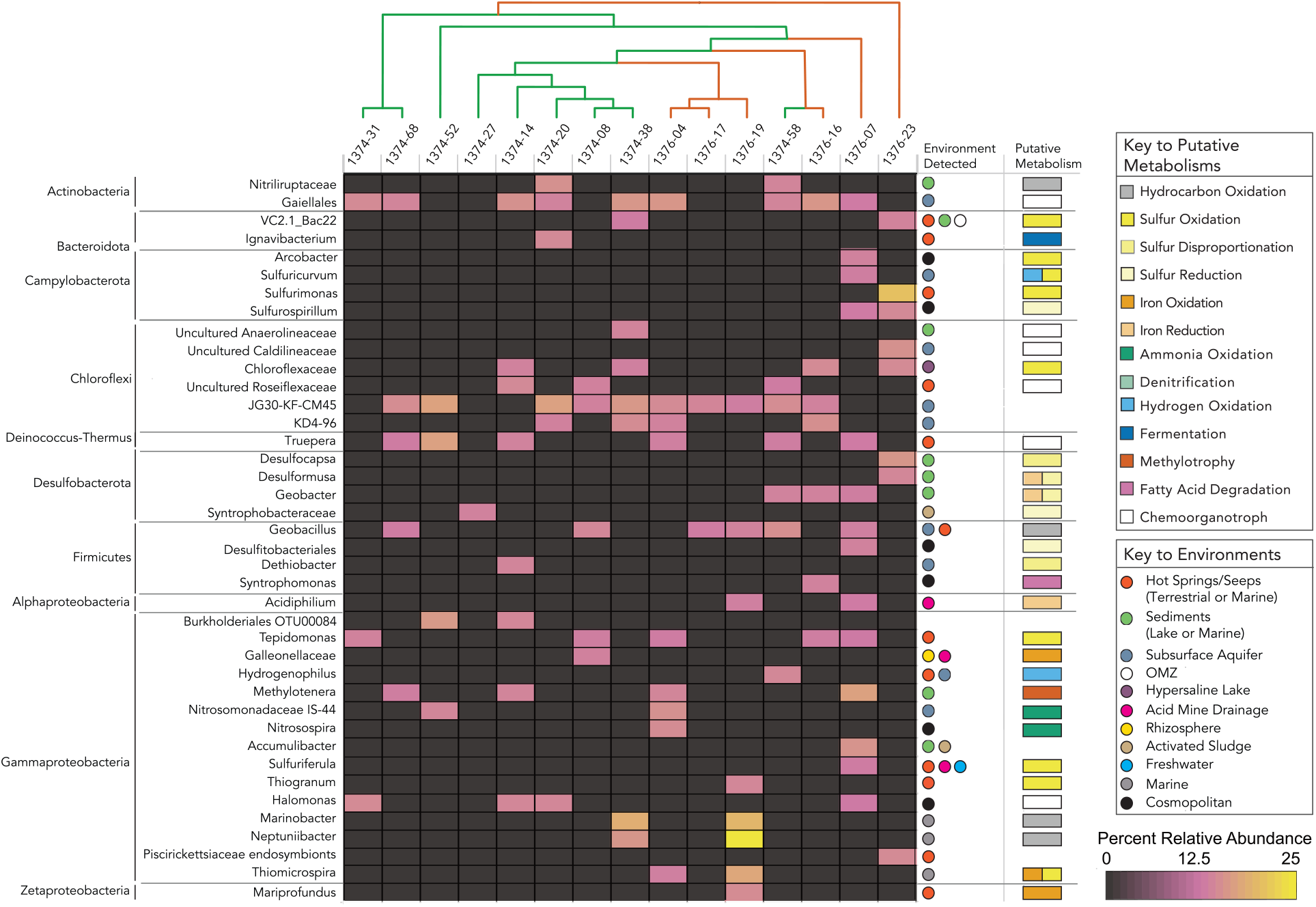
Heatmap of OTUs most likely endemic to the subsurface Louisville Seamounts. Organized by phylum (at left) and lowest classification possible (each row) and sample (columns). A Jaccard dissimilarity cladogram is at top, with samples from Hole U1374A in green and Hole U1376A in red. The environments from which each OTU is typically detected is listed at right, next to the putative metabolism for those OTUs where significant cultured representatives have been studied.

We focused on analysis of genera and families detected that may be indicative of a subsurface lifestyle based on where cultured representatives or environmental sequences have been predominately detected (Figure 4). Many genera and families detected here are also detected in other subsurface settings, including hot springs, cold seeps, sediments and subsurface aquifers^28,48–72^. Of these, there was a predominance of putative metabolisms related to S, N and Fe redox chemistry and hydrocarbon oxidation. The detection of metabolisms related to S and Fe redox chemistry is notable because basalts are composed of ∼1% S and 10% Fe, therefore these sources of metabolic energy are inherent in the substrate. In particular, Zetaproteobacteria related to *Mariprofundis ferooxidans*, an obligate Fe-oxidizer^73^, were detected in the Louisville Seamount chain subsurface. Zetaproteobacteria were also detected in the North Pond observatories, where metagenomic and transcriptomic data indicates they are fixing carbon and oxidizing Fe^22,23^. Other putative iron oxidizers in the Louisville Seamount chain subsurface include *Thiomicrospira*, which was detected in the same sample as *M. ferrooxidans,* and an isolate of which was recently shown to oxidize Fe^50^, and *Gallionaceae*. Putative iron reducers include *Geobacter*, *Desulfuromusa* and *Acidiphilium*. Putative sulfur oxidizers and sulfur reducers, were distributed within the Bacteroidota, Campylobacterota, Desulfobacterota, Firmicutes and Gammaproteobacteria (Figure 4).

Several genera putatively capable of hydrocarbon oxidation were also detected, including *Nitriliruptaceae*, *Geobacillus*, *Thauera*, *Marinobacter* and *Neptuniibacter*. The latter two, in particular, have been noted for their role in hydrocarbon oxidation in the marine environment^53,74,75^. Subsurface communities detected in gabbro below Atlantis Massif and Atlantis Bank included abundant hydrocarbon degraders^26,43^, indicating that this may be a common subsurface phenotype.

To determine if microorganisms from subsurface seamount environments could be stimulated to grow, we initiated enrichment experiments with core samples from Holes U1372A, U1373A, U1374A and U1376A during Expedition 330 and extracted DNA from those experiments 171-1281 days later (Table S3). We used growth media targeting chemoorganotrophs, iron reducers and sulfur oxidizers^76^, likely metabolisms based on the presence of S and Fe in basalts. Analysis of V4V6 16S rRNA amplicons from the 35 enrichment incubations yielded additional insight beyond the amplicon analysis of the cored samples (Fig. S3). In particular, several taxa were recovered in enrichments as well as from the core samples, including the VC2.1_Bac22 clade of Bacteroidia, which is associated with sulfidic environments and capable of sulfur oxidation^47,77^, *Sulfurimonas* (Campylobacterota), *Methylotenera, Thiomicrospira* and *Halomonas* (Gammaproteobacteria) and *Mariprofundus* (Zetaproteobacteria; Figure 5). In total, five Zetaproteobacterial OTUs were detected (Table S4), including ZetaOTU9, considered a subsurface clade^78^ and most closely related to the recently described *Ghiorsea bivora*, which can oxidize both Fe and H^79^; ZetaOTU11, represented by *M. ferrooxidans*; and ZetaOTU58, recently recovered from mild steel incubated in sediments^50,80^. Additionally, the taxa Sphingobacteriaceae (Bacteroidia), the Ktedonobacteria taxa B12-WMSP1 and the genus *Gemmatimonas* in the class Gemmatimonadota were found exclusively in enrichments with added Fe(III), indicating a possible role in Fe-cycling. Cultivated members of *Thiomicrospira*, *Zetaproteobacteria* and *Sulfurimonas*^50,52^ are autotrophic and therefore the amplicons detected here putatively represent members of the base of the subsurface food web in the Louisville ecosystem.

**Figure 5.**
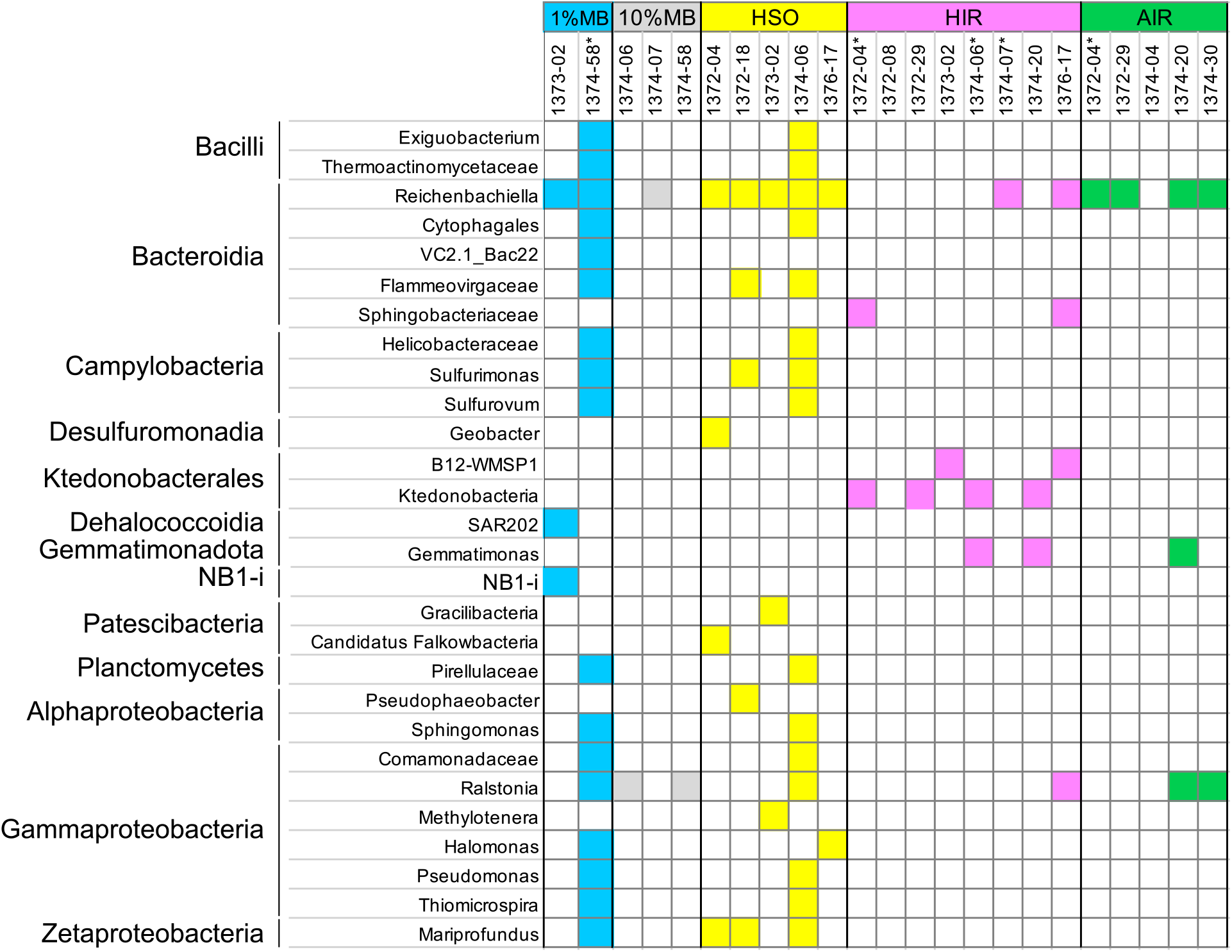
Presence of taxa in enrichment experiments from IODP Expedition 330. Media indicated at the top of the chart, each column represents one sample. 1%MB = 1% Marine Broth in artificial seawater; 10%MB = 10% Marine Broth in artificial seawater, HSO = heterotrophic sulfur oxidizer media; HIR = heterotrophic iron reducer media; AIR = autotrophic iron reducer media. Sample names are the site number followed the core number (e.g. 1374-07 for sample U1374A-7R1). Asterisk indicates two enrichment vials were initiated from the same sample.

Difficulty in sampling and low biomass has previously hampered analysis of the deep subsurface biosphere in subseafloor igneous basement. Knowledge to date is derived largely from the JdFR flank, a warm (∼65°C), hydrothermally active crustal biome^9,12–15^, and North Pond, a young, hydrologically active ridge flank with cool (∼4-20°C), oxic waters^20,21,24,25^, but it was thought that sites older than 65 Ma should be inactive due to crustal sealing^30–32^. We showed here that there is indeed a subseafloor crustal biome in these older regions of seafloor that is unique from that in the younger setting. This is important because it reveals that the subseafloor crustal biome cannot be treated as a monolithic entity as we endeavor to understand global biogeochemical processes. It also indicates that considerations of global subsurface biomass must include marine basement of all ages, which will impact these estimates given the massive size of the habitat. Further, the biomass values measured here, which included data from the JdFR rocks in addition to the Louisville Seamount rocks, in addition to previous work at North Pond^21,24,25^, Atlantis Massif^27^ and Atlantis Bank^28^, indicate a subseafloor biome with lower cell densities than overlying sediments. However, subseafloor basement is a spatially vast biosphere, including hotspots of biomass at mid-ocean ridges and upper oceanic crust^36^, so their population size, geographical distribution, metabolic activity, and ecological significance remain largely unknown. Future work will help build a global database of subseafloor microbiomes in this least-explored crustal biosphere.

## Materials and Methods

### Core Handling and Sampling

All samples were collected using rotary core barrel drilling during IODP Expedition 330, 13 December 2010 - 11 February 2011 (Figure 1). Detailed sampling methods are published elsewhere^40^ and summarized here. Whole-round cores were selected in the core splitting room and collected from the core liner onto pre-combusted (450°C for 2 hours) aluminum foil. Sections were specifically chosen that showed some sign of alteration or a fluid flow conduit because these are likely locations for microbial life. Microbiology samples ranged 5-14 cm long. Prior work has shown that the interior of rock cores is generally free from contamination^81^, therefore, efforts were taken to sample only the interior of the cores. The intact whole round was washed 3X with artificial seawater in a fresh ziplock bag for each rinse before subsampling to avoid contamination from drilling fluids. Next, the rock was split with a flame-sterilized sterile chisel and sampled for the interiors of the cores. Samples were placed in 5 mL, autoclaved centrifuge tubes and immediately put at - 80°C for later analysis. Details of contamination testing is supplied in the Supporting Online Materials.

### Biomass enumeration

A cell extraction and enumeration method originally developed for quantifying microbial biomass in subseafloor sediments^82^ and recently adapted for samples from ocean crust^27,28^ was used. Briefly, frozen samples were powderized in a tungsten carbide mortar and pestle that was previously decontaminated with RNAse Away, and then fixed in sterile filtered 2% formaldehyde at a volume:volume ratio of 1:5. One ml of the fixed sample slurry was used in the quantification procedure, which was followed as in Morono et al. (2013) except that 40 cycles of sonication were used instead of 20 to liberate cells attached to the powdered rock. Cells were enumerated on filters stained with 1/40 SYBR Green 1 in TE buffer by counting either 800-900 fields of view if fewer than 40 cells total were detected, or at least 40-50 cells in fewer fields when possible. The limit of quantification was defined as 3X the standard deviation of the mean of the negative control counts. Two nwegative controls were processed and analyzed for every ten experimental samples.

### DNA extraction and sequencing from cores

Core samples for both amplicon and metagenomic analysis were extracted as previously^47^ using a CTAB phenol/chloroform protocol with 1% CTAB. Starting material for amplicon sequencing samples was ∼4 cm^3^ of rock chips. With the exception of sample 1376-23, all other samples analyzed by amplicon analysis were purified using synchronous coefficient of drag alteration (SCODA)^83^, as implemented with disposable cartridges and the Aurora System (Boreal Genomics, Mountain View, CA). A SCODA negative control was processed by running ultrapure deionized water in an Aurora cartridge and sequencing the output.

### Enrichment experiments

Enrichment experiments were started during Expedition 330 (see Table S3 for details). For each, ∼1 cm^3^ was added to a serum vial with 5 ml media. The media used targeted heterotrophs (1% Marine Broth and 10% Marine Broth), heterotrophic sulfur oxidizers (HSO), autotrophic iron reducers (AIR) or heterotrophic iron reducers (HIR) using recipes published previously^76^. All enrichments were incubated at 4°C until processed for DNA extractions. Five ml of enrichment culture was filtered onto a 0.2 μm pore-size polycarbonate filter and frozen at -80°C until community DNA was extracted using either the same method as used for the core samples (see main text for details), MoBio PowerWater DNA Isolation Kit, or MP Biomedical FastDNA Kit for Soil, as indicated in Table S3. Samples were sent to Research and Testing Lab for PCR of the V4V6 region of 16S rRNA using the same primers as the core samples and then sequencing using 454 pyrosequencing with a target of 3000 reads per sample.

One sample, 1372-18-HSO, was processed using the bacterial primers 27F (5’- GAG TTT GAT CCT GGC TCA G-3’) and 519R (5’-GTA TTA CCG CGG CTG CTG G-3’) for PCR followed by cloning and sequencing of the PCR products. The PCR product was run on an agarose gel, cut out, and extracted using QIAquick Gel Extraction Kit (Qiagen, Valencia, CA, USA) according to the manufacturer’s instructions. Fragments were cloned into the pCR 4 TOPO vector using the TOPO TA Cloning Kit (Invitrogen, Grand Island, NY, USA) and transformants plated on LB+100 μg mL^-1^ ampicillin according to the manufacturer’s instructions. Colonies were randomly selected and grown in liquid culture followed by sequencing at Beckman Coulter Genomics (Danvers, MA).

The 454 amplicon datasets were processed in mothur^84^ using the same protocol as for the core samples. Clones were assessed via BLAST to determine identity at the genus level; only five unique sequences were recovered. All data is available in the NCBI Sequence Read Archive under Bioproject PRJNA271884, accession numbers SRX834467-SRX834482 (core samples), Bioproject PRJNA1033664, accession numbers SAMN38041014-SAMN38041051 (enrichment samples), and Genbank accession numbers OR751581-OR751585 (representative clones from sample 1372-18-HSO).

### Archaea specific quantitative Polymerase Chain Reaction (qPCR)

qPCR was used to estimate relative abundance of Archaea as described previously ^47^ using the primers 806f (5’-ATT AGA TAC CCS BGT AGT-3’ ^85^) and 922r (5’-YCC GGC GTT GAN TCC AAT T-3’ ^86^). The thermal program employed was: 10 minutes at 95°C followed by 45 cycles of 30 sec. at 95°C, 30 sec. at 55°C and 25 sec. at 72°C. Melt curves for all qPCR products were checked to ensure a single PCR product was generated. qPCR reactions were run in triplicate and the limit of detection was 54 gene copies per reaction, the mean of qPCR negative control reactions with water added instead of sample.

### 16S rRNA Amplicon Analysis

V4V6 amplicons were analyzed using the software package mothur ^84^ in two rounds, adapting elements of a protocol used to determine contamination in deep subsurface sediment samples ^87^. OTUs have been found to yield similar ecological results to amplicon sequence variants ^88^, including in deep subsurface biosphere environments ^89^, and were used here. For the first round, OTUs were generated at the 95% cutoff level. All OTUs were generated using the pre.cluster option in Mothur, which uses modified single-linkage ^90^ to account for sequencing errors. Four mismatches were allowed per cluster, equivalent to one mismatch per 100 bp ^91^ and the average neighbor method was used for OTU clustering. To control from contamination, OTUs detected in either the negative control samples (73DF or AuroraBlank) at an average relative abundance of ≥10X than they were detected in the average of the 15 experimental samples were removed. Additionally, taxa were removed that were previously detected in sequencing kits as contaminants (^92^, Table S5). The contaminant subtracted amplicon libraries from the 15 experimental samples were then run through the same analysis pipeline in mothur starting from the raw sequences, this time using 97% similarity cutoff to generate OTUs for comparison of OTU abundance per sample and 95% for comparison of community similarity between samples, as has been done previously ^47,93^.

## Acknowledgements

This paper is dedicated to the memory of Katrina Edwards, who was a life-changing mentor to JBS and a pioneer in the field of subseafloor microbiology. The authors would like to thank IODP Expedition 330 Co-Chief Scientist Toshitsugu Yamazaki as well as Staff Scientist Joerg Geldmacher. We thank Jordan Hoese and Arik Joukhajian for assistance analyzing enrichment experiments. This is a contribution to the Deep Carbon Observatory (DCO).

