## Supplementary Online Material for "Microbial abundance and diversity in 64-74 Ma subseafloor igneous basement from the Louisville Seamount Chain"

#Deceased

### Supplementary Methods

*Contamination testing*

As part of the drilling process, huge amounts of surface seawater mixed with a proprietary chemical mix are injected into boreholes; this fluid, known as drilling fluid or drilling mud, is the major source of contamination of microorganisms in cores collected during coring. To check for contamination, the microbial composition of drilling fluid was assessed (Sample 73DF). 73DF was collected directly from the injection pipe on deck into sterile bottles with screw caps and then frozen at -80°C until processing. Once thawed, microorganisms present in fluids were collected by filtration using a vacuum pump onto a 0.2 µm pore polycarbonate filter. DNA was extracted as described below immediately following filtration.

Bags of yellow to green fluorescent microspheres (Fluoresbrite carboxylate microspheres; Polysciences Inc. 15700) with a diameter of 0.52 (±0.01) µm were used as a particulate tracer that mimics microbial cells on the *JOIDES Resolution* [1]. These microspheres were counted in core samples for an indication of contamination on 2-3 cores per site.  The concentration of microspheres was set at $10^{10}$ spheres/ml[1]. Concentrations of fluorescent microspheres in core samples were quantified using a Zeiss Axiophot epifluorescence microscope outfitted with a mercury lamp (HBO 100W), a blue filter set, and a 100Å~ Plan-NEOFLUAR oil-immersion objective.  Non-fluorescent immersion oil was used for all observations.  Aliquots (100µL) of the crushed rock were suspended into 10 ml of filtered 1xPBS solution and filtered onto black, 25-mm-diameter polycarbonate filters (0.2-µm pore size) in a filtration tower. The microspheres on the filter were then counted using the epifluorescence microscope.  Microsphere abundance on the filters was determined by averaging the total number seen in at least 20 randomly selected fields of view and by looking at outer, inner and center portions of the core.

#### Supplemental Results

*Discussion of qPCR results*

qPCR was performed on samples after they were cleaned up with the Boreal Genomics Aurora Purification System, as described in the Materials and Methods section.  Bacterial 16S rRNA was quantified in triplicate as described previously[2] using primers 338f (5′-ACT CCT ACG GGA GGC AGC AG-3') and 518r (5'-ATT ACC GCG GCT GCT GG-3'.  qPCR for Archaea was carried out as described in Materials and

Methods. Because there was obvious background contamination in the Aurora cartridges, we decided the data for bacterial biomass is unreliable. However, because archaeal 16S rRNA was below detection, we believe this reflects the environment accurately because the contamination should result in either similar or increased 16S rRNA gene copies. Dilution below the detection limit is also possible, but it should be noted that other similar environments also have low abundance of Archaea[3,4]. Therefore, the lack of archaeal amplification is reported as part of our analysis since it indicates a true lack of archaeal 16S rRNA gene copies, but the bacterial quantification is not reported as it will clearly be biased.

#### Supplementary Online Material References


1. Smith DC, Spivack AJ, Fisk MR, Haveman SA, Staudigel H, Party tLSS. Methods for quantifying potential microbial contamination during deep ocean coring. ODP Technical Note. 2000;28.
2. Einen J, Thorseth IH, Ovreas L. Enumeration of Archaea and Bacteria in seafloor basalt using real-time quantitative PCR and fluorescence microscopy. FEMS Microbiol Lett. 2008;282(2):182-7.
3. Jorgensen SL, Zhao R. Microbial inventory of deeply buried oceanic crust from a young ridge flank. Frontiers in Microbiology. 2016;7:820.
4. Wee SY, Edgcomb VP, Beaudoin D, Yvon-Lewis S, Sylvan JB. Microbial Abundance and Diversity in Subsurface Lower Oceanic Crust at Atlantis Bank, Southwest Indian Ridge. Applied and Environmental Microbiology. 2021;87(22):e0151921.

### Supplementary Online Tables

Table S1 - Summary of $\delta^{13}C$-Total C, Total C content, $\delta^{13}C$-TOC and TOC content

| Sample | Depth Below Seafloor (mbsf) | Cell Biomass (cells $cm^{-3}$) | Total C $\delta^{13}C$ ‰VPDB | Total C (wt %) | TOC $\delta^{13}C$ ‰VPDB | TOC (wt %) |
|---|---|---|---|---|---|---|
| U1374A-07-R-1 | 35.4 | 8334 | 2.02 | 0.377 | -20.36 (n=2) | 0.007 (n=2) |
| U1374A-20-R-1 | 102.9 | 247 | 2.67 | 3.231 | -20.54 (n=2) | 0.013 (n=2) |
| U1374A-27-R-1 | 135.8 | 168 | 2.74 | 3.465 | -21.67 | 0.010 |
| U1374A-38-R-2 | 203.6 | 292 | 2.85 | 1.535 | -20.11 | 0.008 |
| U1374A-58-R-6 | 400.2 | 716 | -6.99 (n=2) | 0.142 (n=2) | -17.83 | 0.014 |
| U1376A-04-R-1 | 29.4 | 1443 | 3.77 | 11.776 | -23.68 | 0.154 |
| U1376A-17-R-2 | 126.7 | 124 | -1.59 | 0.767 (n=2) | -23.048 (n=2) | 0.006 (n=2) |
| U1376A-19-R-1 | 144.6 | bql | 0.46 | 1.703 | | |

Table S2 - Diversity statistics for bacterial pyrotags

| Short Sample Name | # OTUs | Chao 1 | ACE |
|---|---|---|---|
| 1374-08 | 175 | 307 | 236 |
| 1374-14 | 232 | 317 | 281 |
| 1374-20 | 180 | 271 | 224 |
| 1374-27 | 149 | 317 | 236 |
| 1374-31 | 163 | 913 | 1786 |
| 1374-38 | 218 | 323 | 289 |
| 1374-52 | 100 | 408 | 698 |
| 1374-58 | 271 | 715 | 859 |
| 1374-68 | 138 | 610 | 685 |
| 1376-04 | 275 | 803 | 743 |
| 1376-07 | 325 | 919 | 1605 |
| 1376-16 | 260 | 837 | 875 |
| 1376-17 | 257 | 635 | 817 |
| 1376-19 | 173 | 433 | 615 |
| 1376-23 | 209 | 510 | 592 |

Table S3 - Sample data for enrichment incubations.

| Sample | Depth (mbsf) | Date Sampled on Exp330 | Transfer Date | Days Incubated | DNA Extraction Method | # Pyrotags (post-subtraction) |
|---|---|---|---|---|---|---|
| U1373A-2R1-1MB | 10.8 | 1-Jan-11 | 17-Aug-11 | 228 | MoBio PowerWater | 2386 (2386) |
| U1374A-58R6-1MB | 400.2 | 15-Jan-11 | 5-Jul-11 | 171 | MoBio PowerWater | 1899 (1175) |
| U1374A-6R3-10MB | 33.4 | 7-Jan-11 | 7-Nov-13 | 1035 | MP Biomedical Fast Soil | 3280 (3280) |
| U1374A-7R1-10MB | 35.4 | 7-Jan-11 | 25-Jan-13 | 749 | MP Biomedical Fast Soil | 1397 (1389) |
| U1374A-7R1-10MB | 35.4 | 7-Jan-11 | 7-Nov-13 | 1035 | MP Biomedical Fast Soil | 1462 (1462) |
| U1372A-4R3-HSO | 16.9 | 22-Dec-10 | 17-Aug-11 | 238 | MoBio PowerWater | 1420 (1418) |
| U1372A-18R2-HSO | 135.3 | 25-Dec-10 | 17-Aug-11 | 235 | MoBio PowerWater | NA |
| U1373A-2R1-HSO | 10.8 | 1-Jan-11 | 17-Aug-11 | 228 | MoBio PowerWater | 1293 (1293) |
| U1374A-6R3-HSO | 33.4 | 7-Jan-11 | 12-Sep-11 | 248 | CTAB Phenol/Chloro | 6392 (3995) |
| U1376A-17R2-HSO | 127.8 | 1-Feb-11 | 25-Aug-11 | 205 | CTAB Phenol/Chloro | 1688 (1688) |
| U1372A-4R3-HIR-1 | 16.9 | 22-Dec-10 | 23-Jun-14 | 1279 | MP Biomedical Fast Soil | 6258 (166) |
| U1372A-4R3-HIR-2 | 16.9 | 22-Dec-10 | 25-Jun-14 | 1281 | MP Biomedical Fast Soil | 4620 (4) |
| U1372A-8R5-HIR | 48.0 | 23-Dec-10 | 25-Jun-14 | 1280 | MP Biomedical Fast Soil | 3467 (93) |
| U1372A-18R2-HIR | 135.3 | 25-Dec-10 | 23-Jun-14 | 1276 | MP Biomedical Fast Soil | 2023 (10) |
| U1372A-29R1-HIR-1 | 186.2 | 27-Dec-10 | 25-Jun-14 | 1276 | MP Biomedical Fast Soil | 2674 (4) |
| U1373A-2R1-HIR-1 | 10.8 | 1-Jan-11 | 25-Jun-14 | 1271 | MP Biomedical Fast Soil | 3116 (7) |
| U1374A-4R1-HIR-1 | 21.5 | 6-Jan-11 | 24-Jun-14 | 1265 | MP Biomedical Fast Soil | 2568 (3) |
| U1374A-4R1-HIR-2 | 21.5 | 6-Jan-11 | 25-Jun-14 | 1266 | MP Biomedical Fast Soil | 3136 (93) |
| U1374A-6R3-HIR-1 | 33.4 | 7-Jan-11 | 23-Jun-14 | 1263 | MP Biomedical Fast Soil | 747 (25) |
| U1374A-6R3-HIR-2 | 33.4 | 7-Jan-11 | 30-Jun-14 | 1270 | MP Biomedical Fast Soil | 364 (6) |
| U1374A-7R1-HIR-1 | 35.4 | 7-Jan-11 | 24-Jun-14 | 1264 | MP Biomedical Fast Soil | 6231 (15) |
| U1374A-7R1-HIR-2 | 35.4 | 7-Jan-11 | 25-Jun-14 | 1265 | MP Biomedical Fast Soil | 4177 (39) |
| U1374A-20R1-HIR | 102.9 | 8-Jan-11 | 25-Jun-14 | 1264 | MP Biomedical Fast Soil | 5554 (18) |
| U1374A-30R3-HIR | 156.8 | 10-Jan-11 | 25-Jun-14 | 1262 | MP Biomedical Fast Soil | 4938 (2) |
| U1376A-16R5-HIR | 120.8 | 31-Jan-11 | 23-Jun-14 | 1239 | MP Biomedical Fast Soil | 8512 (5) |

| | | | | | | |
|---|---|---|---|---|---|---|
| U1376A-17R2-HIR | 127.8 | 1-Feb-11 | 23-Jun-14 | 1238 | MP Biomedical Fast Soil | 5379 (44) |
| U1376A-23R1-HIR | 174.2 | 2-Feb-11 | 25-Jun-14 | 1239 | MP Biomedical Fast Soil | 6701 (8) |
| U1372A-4R3-AIR-1 | 16.9 | 22-Dec-10 | 30-Jun-14 | 1286 | MP Biomedical Fast Soil | 1205 (21) |
| U1372A-4R3-AIR-2 | 16.9 | 22-Dec-10 | 30-Jun-14 | 1286 | MP Biomedical Fast Soil | 4454 (4143) |
| U1372A-29R1-AIR-2 | 186.2 | 27-Dec-10 | 30-Jun-14 | 1281 | MP Biomedical Fast Soil | 1484 (21) |
| U1374A-4R1-AIR-2 | 21.5 | 6-Jan-11 | 2-Jul-14 | 1273 | MP Biomedical Fast Soil | 2003 (141) |
| U1374A-7R1-AIR-1 | 35.4 | 7-Jan-11 | 2-Jul-14 | 1272 | MP Biomedical Fast Soil | 571 (8) |
| U1376A-16R5-AIR | 120.8 | 31-Jan-11 | 7-Jul-14 | 1253 | MP Biomedical Fast Soil | 5281 (3) |
| U1374A-20R1-AIR | 102.9 | 8-Jan-11 | 30-Jun-14 | 1269 | MP Biomedical Fast Soil | 2349 (63) |
| U1374A-30R3-AIR-1 | 156.8 | 10-Jan-11 | 30-Jun-14 | 1267 | MP Biomedical Fast Soil | 3153 (15) |
| Extraction Blank | NA | NA | NA | NA | MP Biomedical Fast Soil | 73 (0) |

Table S4 - Assignments of OTUs defined as Zetaproteobacteria to Zetaproteobacterial OTUs (ZetaOTUs), as defined by Zetahunter (McAllister et al., 2018).

| OTU name | Sample | ZetaOTU |
|---|---|---|
| LouiC2_E7 | U1372-18R2-HSO | ZetaOtu9 |
| LouiC2_H10 | U1372-18R2-HSO | ZetaOtu9 |
| Lou10_II8GYES02HQNI9_Otu270 | U1374-58R6-1MB | ZetaOtu9 |
| Lou11_IJDV81G03GGH4L_Otu029 | U1374-6R3-HSO | ZetaOtu9 |
| Lou11_IJDV81G03FXDQE_Otu754 | U1374-6R3-HSO | ZetaOtu9 |
| Lou11_IJDV81G03GAEBC_Otu907 | U1374-6R3-HSO | ZetaOtu9 |
| Otu00110 | U1376-19R1 | ZetaOtu11 |
| Lou10_II8GYES02JGD7D_Otu134 | U1374-58R6-1MB | ZetaOtu58 |
| Lou10_II8GYES02GW1OC_Otu908 | U1374-58R6-1MB | ZetaOtu58 |
| Lou10_II8GYES02GU5GW_Otu906 | U1374-58R6-1MB | ZetaOtu58 |
| Lou10_II8GYES02HBNCU_Otu955 | U1374-58R6-1MB | ZetaOtu58 |
| LouiC2_D6 | U1373-4R3-HSO | ZetaOtu58 |
| Lou11_IJDV81G03G1PT5_Otu011 | U1374-6R3-HSO | ZetaOtu58 |
| Lou11_IJDV81G03HBVVZ_Otu910 | U1374-6R3-HSO | ZetaOtu58 |
| Lou11_IJDV81G03GZGB1_Otu422 | U1374-6R3-HSO | NewZetaOtu1 |
| Lou11_IJDV81G03F7296_Otu390 | U1374-6R3-HSO | NewZetaOtu2 |

#### Supplementary Online Figures

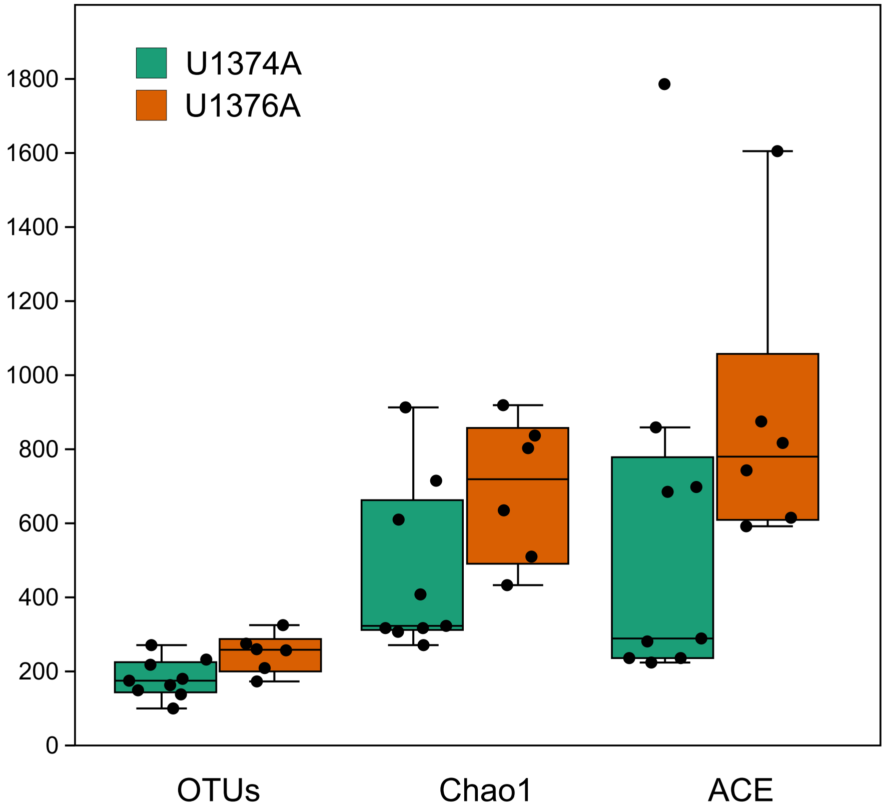


Figure S1 - Box and whisker plots for number of bacterial OTUs, defined at the 97% similarity cutoff, Chao1 and ACE for samples from Holes U1374A and U1376A.

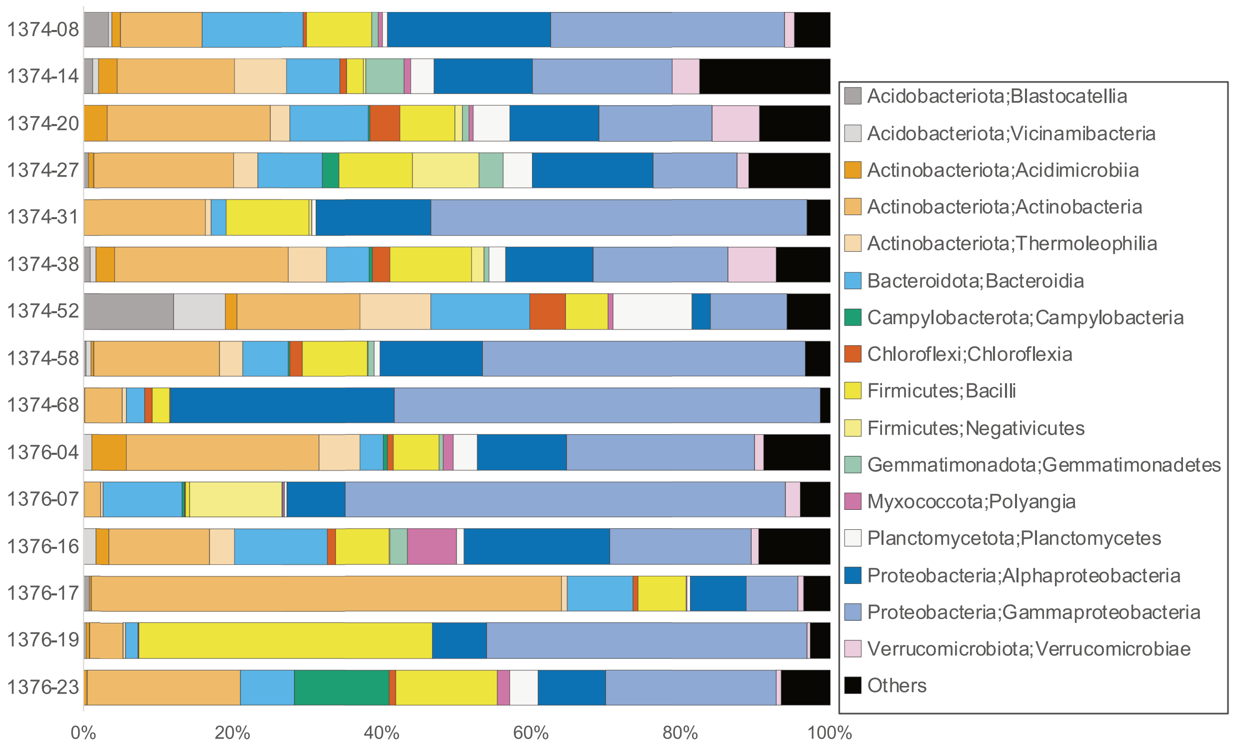


Figure S2 – Bacterial classes detected through amplicon sequencing of the V4V6 region of 16S rRNA. "Others" represents taxa not present at >1% in any sample. Numbers of amplicons per sample are indicated in Table 1.

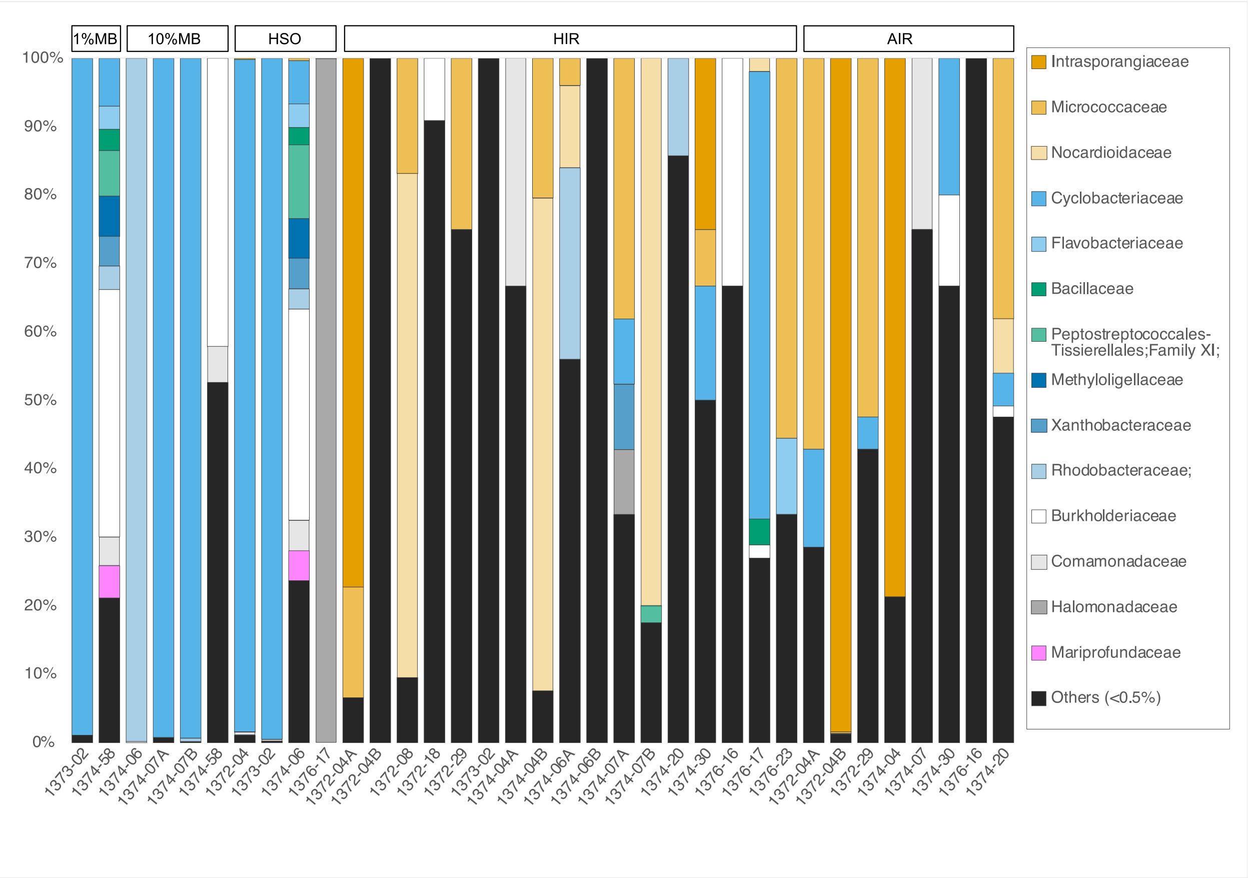


Fig. S3 - Bacterial families detected in enrichment incubations through pyrotag sequencing of the V4V6 region of 16S rRNA. "Others" represents taxa not present at >0.5% in any sample. Numbers of pyrotags per sample are indicated in Table S5. Sample names are the site number followed the core number (e.g. 1374-07 for sample U1374A-7R1).
